# Frequency-coded patterns of sympathetic vasomotor activity are differentially evoked by the paraventricular nucleus of the hypothalamus in the Goldblatt hypertension model

**DOI:** 10.1101/2023.03.13.532381

**Authors:** Jean Faber, Maycon I. O. Milanez, Cristiano S. Simões, Ruy R. Campos

**Affiliations:** Cardiovascular Division, Department of Physiology, Universidade Federal de São Paulo, Escola Paulista de Medicina; Neuroscience Division, Department of Neurology and Neurosurgery, Universidade Federal de São Paulo, Escola Paulista de Medicina

**Keywords:** neural frequency code, hypertension, paraventricular nucleus of the hypothalamus, renal sympathetic activity, splanchnic sympathetic activity, cluster analysis, losartan, bicuculline

## Abstract

The activation of specific brain areas involved in regulating the vasomotor sympathetic activity can lead to distinct effects in the postganglionic nerves in both physiological and pathological conditions, suggesting that the sympathetic vasomotor activity is differentially coded depending on the nerve outflow and the target organs. Previous studies investigating such patterns have mostly focused on the global energy of the signal. However, recent evidence has suggested that relevant information is coded in the power distribution along the frequency range. Disturbing the sympathoexcitatory vasomotor tone in the paraventricular nucleus of the hypothalamus (PVN) allows to investigate the sympathetic nerve activity in overloaded conditions in both hypertensive and control animals. By disinhibiting the PVN through the microinjection of bicuculline, an antagonist of γ-aminobutyric acid type A (GABAa) receptors, in the Goldblatt (2K1C) rat model of hypertension we addressed the territorially differential changes in the frequency parameters of the renal and splanchnic sympathetic nerve activity (rSNA and sSNA, respectively). We also tested the effect of the systemic administration of losartan, an antagonist of the angiotensin II type 1 receptors (AT1), in the attenuation of the increased rSNA and sSNA in 2K1C rats, once these changes are reported to be dependent on the AT1 activation in the Goldblatt model. Our results revealed that each nerve activity presents its own electrophysiological pattern of frequency-coded rhythm in each group, in basal condition and after bicuculline microinjection, but with no significant differences regarding total power comparison among groups. Additionally, the 2K1C animals treated with losartan showed no decrease in the hypertensive response triggered by the GABAa antagonism when compared to the non-treated 2K1C group. However, their spectral patterns of sympathetic nerve activity were different from the other two groups, suggesting that the systemic blockade of AT1 receptors does not totally recover the basal levels of neither the autonomic symptoms nor the electrophysiological patterns in the Goldblatt model, but act on their spectral frequency distribution. These results suggest that the differential responses evoked by the PVN were preferentially coded in frequency of vasomotor sympathetic responses, indicating that the PVN distinctly modulated each rhythmic activity.

Financial Support – FAPESP (2019/25295-0)

## 1. INTRODUCTION

The sympathetic nervous system (SNS) plays a fundamental role in maintaining body homeostasis. The sympathetic activation imbalance is frequently associated with pathological conditions and their severity (Carnagarin et al., 2018; Guyenet, 2006). Vasomotor sympathetic overactivation, for instance, affects the cardiovascular system, leading to hypertension, chronic kidney disease and heart failure (Campos et al., 2015; Carillo et al., 2012; Nishihara et al., 2017).

The reciprocal communication between the peripheral organs and structures of the Central Nervous System (CNS) involved in the vasomotor sympathetic activity regulation controls the cardiovascular function, including areas from the brainstem, basal forebrain and spinal cord (Dampney, 1994, 2016). The activation of these areas, however, can lead to distinct and even opposing effects in different territories under both physiological and pathological conditions (Esler et al., 2018; McAllen and May, 1994; Ramchandra et al., 2012; Rumantir et al., 1999; Salo et al., 2006). The pharmacological stimulation of specific locations of the paraventricular nucleus of the hypothalamus (PVN), for example, may simultaneously trigger the reduction of the renal sympathetic nerve activity (rSNA) and the increase of the activity of the splanchnic, cardiac and adrenal nerves (Deering and Coote, 2000). Given such distinct outcomes triggered by the same stimulation, it seems that the PVN activity is differentially coded downstream, depending on the nerve outflow and the target organs.

Previous studies investigating the differential activity patterns in the sympathetic nerves so far have mostly focused on the global energy of the signal, lacking a better understanding of its distribution over the frequency range (Barman, 2020; Wehrwein and Barman, 2014). The technical evolution of both electrophysiological recordings and signal analysis in the last few decades, however, has shown that global energy analysis alone is not enough to link sympathetic nerve activity to the genesis and development of hypertension (Wehrwein and Barman, 2014). Evidence has emerged suggesting that relevant information is coded in the spectral features of the signal (Barman et al., 1992, 1995; Chang et al., 1999; Milanez et al., 2020b). The firing frequency of the discharges recorded in the vasomotor sympathetic nerves reflects both their central generation and entrainment by the pulsatile input from the arterial baroreceptors while their amplitude is related to the number of recruited fibers (Guild et al., 2010). There is considerable evidence that stimuli independently controlled and affected the two parameters, such as inputs from chemoreceptors and baroreceptors (Guild et al., 2010), although their central organization by the CNS is still poorly understood.

Disturbing the sympathoexcitatory tonus in central structures, such as the PVN, is a useful way of investigating the changes in the peripheral symptoms and in the sympathetic nerve activity under overloaded conditions in hypertension models (Allen, 2002; Barman, 2020; Bergamaschi et al., 1995; Martin et al., 1991; Milanez et al., 2020b). The desinhibition of the PVN through local microinjection of the γ-aminobutyric acid (GABA)-ergic antagonist bicuculline triggers both a robust pressor response and changes in the sympathetic vasomotor activity pattern to the renal, splanchnic and lumbar territories, generating low-frequency bursts unsynchronized to the cardiac cycle (Kenney et al., 2001). It has been shown that these changes depend on the activation of Angiotensin II type 1 (AT1) receptors in the spinal cord (Milanez et al., 2021, 2020a, 2018). In the present work, we take advantage of this approach in a Goldblatt two-kidney, one-clip (2K1C) rat hypertension model to advance the understanding of how the PVN controls sympathetic vasomotor activity in normal and hypertensive rats addressing the territorially differential changes in the frequency parameters of the basal and bursting activities of the renal (rSNA) and splanchnic (sSNA) sympathetic nerves in hypertensive and normal rats during disinhibition of PVN by local microinjection of bicuculline. Additionally, we investigate the effects of the systemic administration of losartan, an antagonist of the AT1 receptors, on the induced responses by bicuculline into the PVN and increased sympathetic nerve activity in 2K1C rats.

## 2. METHODS

### 2.1 Animals and ethical approval

All procedures were approved by the Ethics in Research Committee of the Escola Paulista de Medicina – Universidade Federal de São Paulo (protocol no. 8724270715/15) and performed in accordance with the guidelines of the National Institute of Health. Male Wistar rats (250–350 g) were housed in group cages under controlled temperature of 23 °C and 12/12 h light/dark cycle with free access to food and water.

### 2.2 Induction of the Goldblatt renovascular hypertension and Losartan treatment

Five-week-old animals were anesthetized intraperitoneally with ketamine (80 mg/Kg) + xylazine (10 mg/Kg). A silver clip (gap width⍰=⍰0.2⍰mm) was carefully implanted around the left renal artery for the induction of the two-kidney, one-clip (2K1C) Goldblatt renovascular hypertension. Five weeks later, some of these animals were treated with losartan (30 mg/kg/day) delivered by oral gavage for 7 consecutive days, as previously described (Nishi et al., 2013).

### 2.3 Experimental and control groups

Experiments were performed 6 weeks after the implantation of the clip, which corresponds to a temporal peak in the sympathetic neurogenic mechanisms underlying the Goldblatt hypertension in the PVN (Campos et al., 2015). The animals were divided in three groups: Goldblatt hypertension-induced rats (2K1C, n=5); Goldblatt hypertension-induced rats treated with losartan (2K1C+LOS, n=5); and age-matched control rats not submitted to Goldblatt surgical procedure (CTRL, n=5).

### 2.4 Recording of mean arterial pressure (MAP) and heart rate (HR)

All animals were independently submitted to ketamine (80 mg/Kg) + xylazine (10 mg/Kg) anesthesia and had the femoral vein catheterized for direct recording of mean arterial pressure (MAP) and heart rate (HR), that were carried out in conscious rats after surgical recovery (approximately 24⍰h) (PowerLab - ADInstruments, Australia). Average values were obtained by continuously recording for 10⍰minutes before urethane intravenous administration.

### 2.5 Electrophysiological recording of renal and splanchnic sympathetic nerve activity and microinjection procedure

Rats were anesthetized with urethane (1.4 g/kg, i.v.). The left renal and splanchnic nerves were retroperitoneally exposed and placed on bipolar silver electrodes, and once the conditions for electrophysiological recording were established, both the nerve and the electrode were covered with paraffin oil.

The animals were then placed in a stereotaxic apparatus (David Kopf, USA) for the microinjection procedure and electrophysiological recording. The PVN was located 1.8 mm caudal to the bregma, 0.5 mm lateral to the midline and 7.8 mm deep from the dorsal medullary surface (bite bar = 3.6 mm) (Paxinos and Watson, 2006).

The baseline signal (BAS condition) from the activity of both nerves was amplified (gain 20 K, NeuroLog, Digitimer, Welwyn Garden City, Herts, UK), bandpass filtered (100 - 1000 Hz), and recorded for 10 minutes using a PowerLab data acquisition system (ADInstruments, Sydney, Australia) for subsequent analysis.

After basal activity recording, the GABAa receptor antagonist bicuculline (400 pMol in 100 nL) was bilaterally microinjected into the PVN using glass micropipettes with tip diameters of 10–20 μm connected to a nitrogen pressure injector (MicroData Instruments Inc., USA), as previously described (Carillo et al., 2012; Oliveira-Sales et al., 2009). Ten minutes after the bicuculline injection (BIC condition), the activity of both nerves was recorded for 10 more minutes.

The acquired signals were analyzed offline using the appropriate software (Spike Histogram – PowerLab – ADInstruments, Australia). The integrated voltage responses of bursting rSNA and sSNA were expressed in arbitrary units (AU) as the change (Δ) from the baseline (BAS) values obtained immediately before each test.

At the end of the experiments, the background noise of sympathetic nerve activity was determined by hexamethonium bromide administration (30 mg/kg, intravenously) to select postganglionic sympathetic ongoing nerve activity. The precision of the microinjection infusion was confirmed by the administration of Evans Blue (2% in 100 nL) into the area. After all proceedings, rats were euthanized with a lethal dose of urethane.

All experimental procedures were conducted in accordance with the Guide for the Care and Use of Laboratory Animals (8th edition, National Academies Press).

### 2.6 Signal pre-processing

All data sets were down-sampled to 1000 Hz of sampling rate and were processed using an IIR notching comb filter to remove 60 Hz and harmonics. Additionally, a bandpass IIR filter was designed with an order of 12 and bandpass ripple 2 dB of 1 and 200 Hz.

### 2.7 Envelope signals

All recordings were enveloped in order to highlight the slow frequencies related to bursts yielded by the injection of bicuculline (BIC) and the slow virtual frequencies of the baseline (BAS) activities. The upper limit of the enveloped signals was calculated using the Matlab® function by implementing the root-mean-square (RMS) envelopes of all 15 recordings, using a sliding window of 50 samples. Figure 1 shows how the sympathetic recordings were assessed.

**Figure 1.**
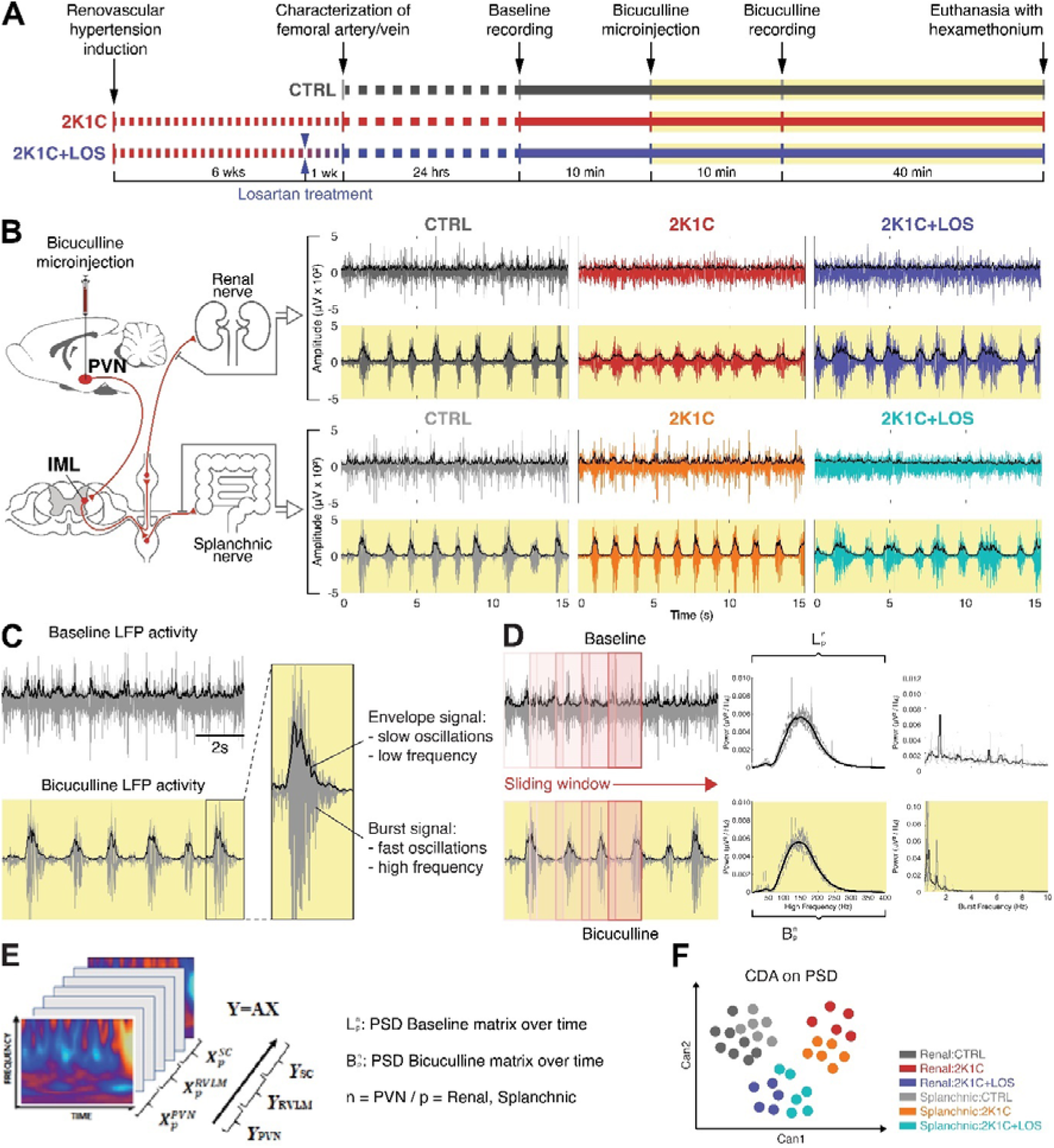
Methods. **(A)** Timeline of the hypertension induction, recordings and bicuculline injection. **(B)** Left: schematic of the projections from the paraventricular nucleus of the hypothalamus (PVN) to the intermediate lateral horn of the spinal cord (IML) and from the postganglionic neurons to the renal and splanchnic territories. Right: recordings of both renal and splanchnic sympathetic nerves activity from baseline (white background) and after bicuculline injection (yellow background). (C) Representative example of the envelope signal at slow oscillations and the filtered signal of burst activity at high oscillations. (D) Schematic of the sliding window analysis and the resulting spectrogram from baseline (white background) and after bicuculline injection (yellow background). (E) Schematic of the generation of the canonical discriminant analysis (CDAs) from the power spectral density (PSDs) distributions. (F) Illustrative example of a scatter plot generated from the Canonical Discriminant Analysis (CDA) with two canonical variables. **(B and F)** Dark gray: Renal nerve, CTRL group; Dark blue: Renal nerve, 2K1C group; Red: Renal nerve, 2K1C+LOS group; Light gray: Splanchnic nerve, CTRL group; Orange: Splanchnic nerve, 2K1C group; Turquoise: Splanchnic nerve, 2K1C+LOS group.

### 2.8 Power spectrum density (PSD)

The (local field potential) LFP recordings were analyzed according to their power spectrum in the frequency domain. The spectrograms were generated to evaluate the power of the frequencies over time, where the Power Spectrum Density (PSD) per animal (n=5 per group, with mean PSD estimated for each group), nerves (renal and splanchnic) and condition (BAS and BIC) were calculated by sliding a temporal window along each recording. The time-window length was heuristically selected, considering 0.5% of the total signal, and 20% of overlap for the sliding steps, in order to maximize temporal and frequency resolutions.

### 2.9 Canonical Discriminant Analysis (CDA)

The canonical discriminant analysis (CDA) was applied on the spectrograms attempting to discriminate the shape of their power density distribution (Guang and Maclean, 2000; Pinheiro et al., 2020; Reyes-Garcia et al., 2018). All the PSDs calculated from each animal and nerve for both conditions (BAS and BIC) were grouped as shown in figure 1F in order to evaluate if they had distinct and characteristic power shapes in specific frequency ranges. Here, we consider that each frequency range corresponds to the set of independent variables. In this way, if the shape of the power density distribution has enough information to discriminate each group described by the categorical dependent variable, the CDA will be able to identify and separate them (Fig. 1G).

To highlight the similarities and differences between the clusters generated by the CDA, we also plotted the arrangement and relationship between the centroids of each cluster. This analysis allowed us to more accurately assess the effect of bicuculline and losartan on the different conditions and nerves.

### 2.10 Statistical analysis

Kolmogorov-Smirnov test was applied to verify the probability of samples coming from a normal distribution. Representations for statistical comparisons were made using mean/median ± CI for parametric and non-parametric data, respectively. To verify statistical differences associated with each nerve and condition, multivariate and bivariate tests were used, such as n-way ANOVA and T-test (or Kruskal-Wallis and Mann-Whitney test for nonparametric distributions).

Kruskal-Wallis test was performed to analyze the differences in power and synchronicity (through interval inter-bursts). For all analyses, a significance level α=5% was used, and all signal processing and statistical analyses were performed using MATLAB® functions with homemade scripts (version 9.2.0 R2018a, Mathworks Inc., MA, USA).

## 3. RESULTS

### 3.1 Systemic treatment with losartan did not revert to the altered parameters in the 2K1C animals after bicuculline injection in the PVN

The systemic administration of losartan (2K1C+LOS group) was not enough to significantly revert the increased response in the mean arterial pressure (MAP, Fig. 2A) and the decreased response in the heart rate (HR, Fig. 2B) between bicuculline and baseline conditions observed in the hypertensive animals (2K1C group). Likewise, it had no significant effect in reducing the over-activation of the renal (rSNA, Fig. 2C) and splanchnic nerves (sSNA, Fig. 2D) after bicuculline injection into the PVN.

**Figure 2.**
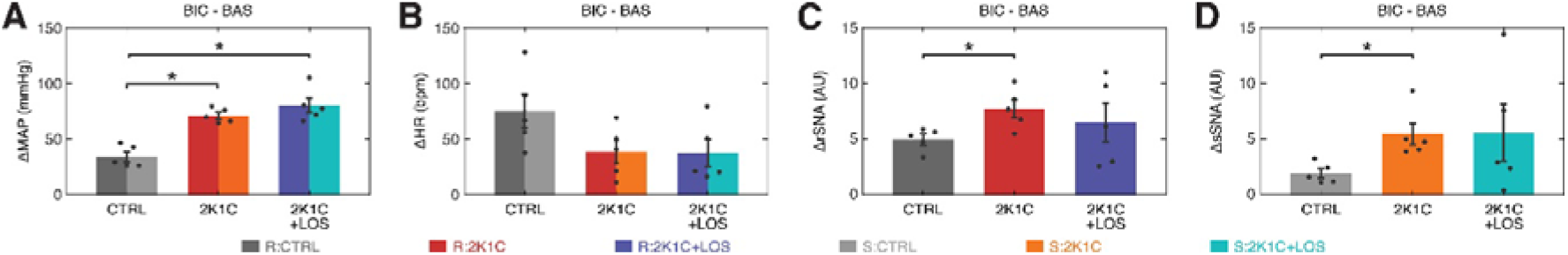
Systemic treatment with Losartan did not revert the altered parameters in the 2K1C animals after bicuculline. **(A)** The increase in the mean arterial pressure (MAP) levels after bicuculline injection from baseline is higher in both hypertensive groups (2K1C and 2K1C+LOS) as compared to the control group (CTRL) regardless of the Losartan treatment. (B) Although not statistically significant, the increase in the heart rate (HR) after bicuculline injection from baseline levels is higher in the CTRL group as compared to both hypertensive groups (2K1C and 2K1C+LOS). **(C, D)** The increase in both renal and splanchnic nerves activity (rSNA and sSNA respectively) after bicuculline injection from baseline levels is significantly higher in the 2K1C group as compared to the CTRL group, but not to the 2K1C+LOS group. **(A, B, C and D)** Values plotted as mean ± SEM. *p<0.05 (One-way ANOVA followed by Bonferroni’s post-hoc test). **(A-D)** Dark gray: Renal nerve, CTRL group; Dark blue: Renal nerve, 2K1C group; Red: Renal nerve, 2K1C+LOS group; Light gray: Splanchnic nerve, CTRL group; Orange: Splanchnic nerve, 2K1C group; Turquoise: Splanchnic nerve, 2K1C+LOS group.

### 3.2 Bicuculline increases the burst envelope power at the 0-4 Hz band with no effect on the high frequency burst activity

At visual inspection, two main frequency bands stand out in the electrophysiological activity of the two nerves during baseline: 0-2 Hz and 6-8 Hz (Fig. 3A and 3B), with the 2K1C+LOS group exhibiting higher power peaks at 5 Hz and 8 Hz than the other groups. After bicuculline administration, these peaks disappear and the power distribution shifts to two main components, one around 0.5 Hz and a weaker one around 1.5 Hz, which is in accordance with previous findings (Kenney et al., 2001) suggesting that the bicuculline injection in the PVN induces a highly synchronized pattern of low frequency activity in both nerves.

**Figure 3.**
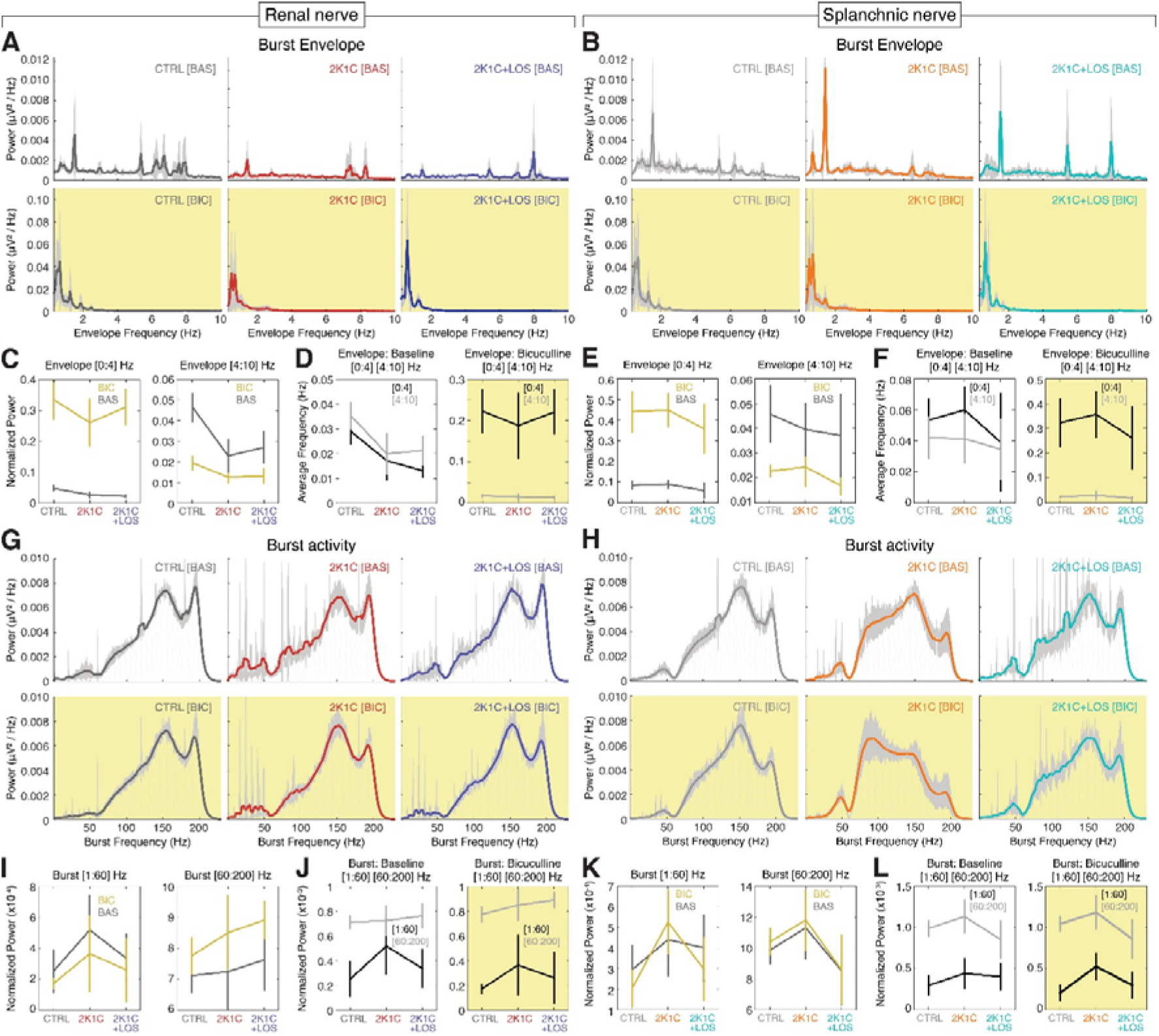
Bicuculline increases the burst envelope power at the 0-4 Hz band and the burst frequency power at the 60-200 Hz band. **(A, B, G, H)** Mean PSD (power spectral density) spectrograms of the low frequency envelopes of the rSNA (A) and sSNA (B) and of the high frequency bursts of the rSNA (G) and sSNA (H) at baseline (top rows, white background) and after bicuculline injection (bottom rows, yellow background). Confidence intervals (CI) values in light gray. **(C, E, I, K)** Mean power of the rSNA (C) and sSNA (E) at the 0-4 Hz (left) and 4-10 Hz (right) frequency bands and of the rSNA (I) and sSNA (K) at the 1-60 Hz (left) and 60-200 Hz (right) frequency bands at baseline (dark gray line) and after bicuculline injection (yellow line). **(D, F, J, L)** Comparison of the average frequency of the maximum power of the rSNA (D) and sSNA (F) between 0-4 Hz (dark gray line) and 4-10 Hz (light gray line) bands at baseline (left) and after bicuculline injection (right) and of the rSNA (J) and sSNA (L) between 1-60 Hz (dark gray line) and 60-200 Hz (light gray line) bands at baseline (left) and after bicuculline injection (right). **(A, B, G and H)** Dark gray: Renal nerve, CTRL group; Dark blue: Renal nerve, 2K1C group; Red: Renal nerve, 2K1C+LOS group; Light gray: Splanchnic nerve, CTRL group; Orange: Splanchnic nerve, 2K1C group; Turquoise: Splanchnic nerve, 2K1C+LOS group.

The bicuculline microinjection increased the bursting activity envelope power of the lower frequencies (0 to 4 Hz) in both nerves and all three groups, without significant differences among groups (Fig. 3C and 3E, left panels). On the other hand, it had the opposite effect on the frequency band between 4 and 10 Hz, although not significant in the 2K1C group in both nerves and in the 2K1C+LOS group in the splanchnic nerve (Fig. 3C and 3E, right panels).

There was no statistical difference between the 0-4 Hz and the 4-10 Hz bands at baseline (Fig. 3D and 3F, left panels). Bicuculline injection, however, promotes a clear separation between the two bands by increasing the 0-4 Hz power and decreasing the 4-10 Hz power (Fig. 3D and 3F, right panels). This statistical effect corroborates the patterns of activity observed in figures 2A and 2B.

With respect to the high frequency bursts, no major differences can be noted at visual inspection among the spectral patterns of the renal and splanchnic nerve activity before and after bicuculline injection. The power distribution is clearly concentrated at frequencies above 60 Hz in both nerves and all three groups, with peaks at around 150 Hz and 190 Hz, which can be seen at a visual inspection of the PSDs (Figs. 2G and 2H). The only exception is the splanchnic nerve activity in the 2K1C group after bicuculline injection that presents a shift in the power peak to around 90 Hz.

No significant differences were observed in the mean power of renal or splanchnic nerve activity in any of the three groups between baseline and bicuculline in the high frequency bursting activity (Fig. 3I and 3K). Comparisons between frequency ranges, however, confirmed that the 60-200 Hz power is higher than the 1-60 Hz in all groups, except for 2K1C in the renal nerve, without significant effect of the bicuculline injection in this pattern (Fig. 3J and 3L).

### 3.3 Clustering analysis using Canonical Discriminant Analysis (CDA) revealed specific frequency patterns on each nerve for each of the three groups (CTRL, 2K1C and 2K1C+LOS) and two conditions (BAS and BIC)

By means of the CDA we performed a joint multivariate analysis considering all the three groups (*CTRL, 2K1C* and *LOS*), both nerves (renal and splanchnic) and each type of activity (baseline and bicuculline administration) for all the five rats simultaneously at the high frequency range of the bursting activity as well as the low range of the envelope oscillations. The advantage of this analysis is that it provides a statistical global view highlighting the homogeneity among animals and heterogeneity among groups.

The cluster analysis was carried out with the first three canonical variables from the CDA using the PSDs shapes as the main feature (Fig. 4). The clusters are formed by the first three canonical variables values plotted against its respective axis in a three dimensional space. Each cluster thus represents the spectral activity distribution across the frequency range for a specific group and nerve. The spectral patterns of all five animals were included, meaning that the statistical effects take into account all three groups (CTRL, 2K1C and 2K1C+LOS) and two nerves (renal and splanchnic) simultaneously for each frequency band (low frequency envelope oscillations, fig. 4A to 4E; high frequency bursts, fig. 4F to 4J).

**Figure 4.**
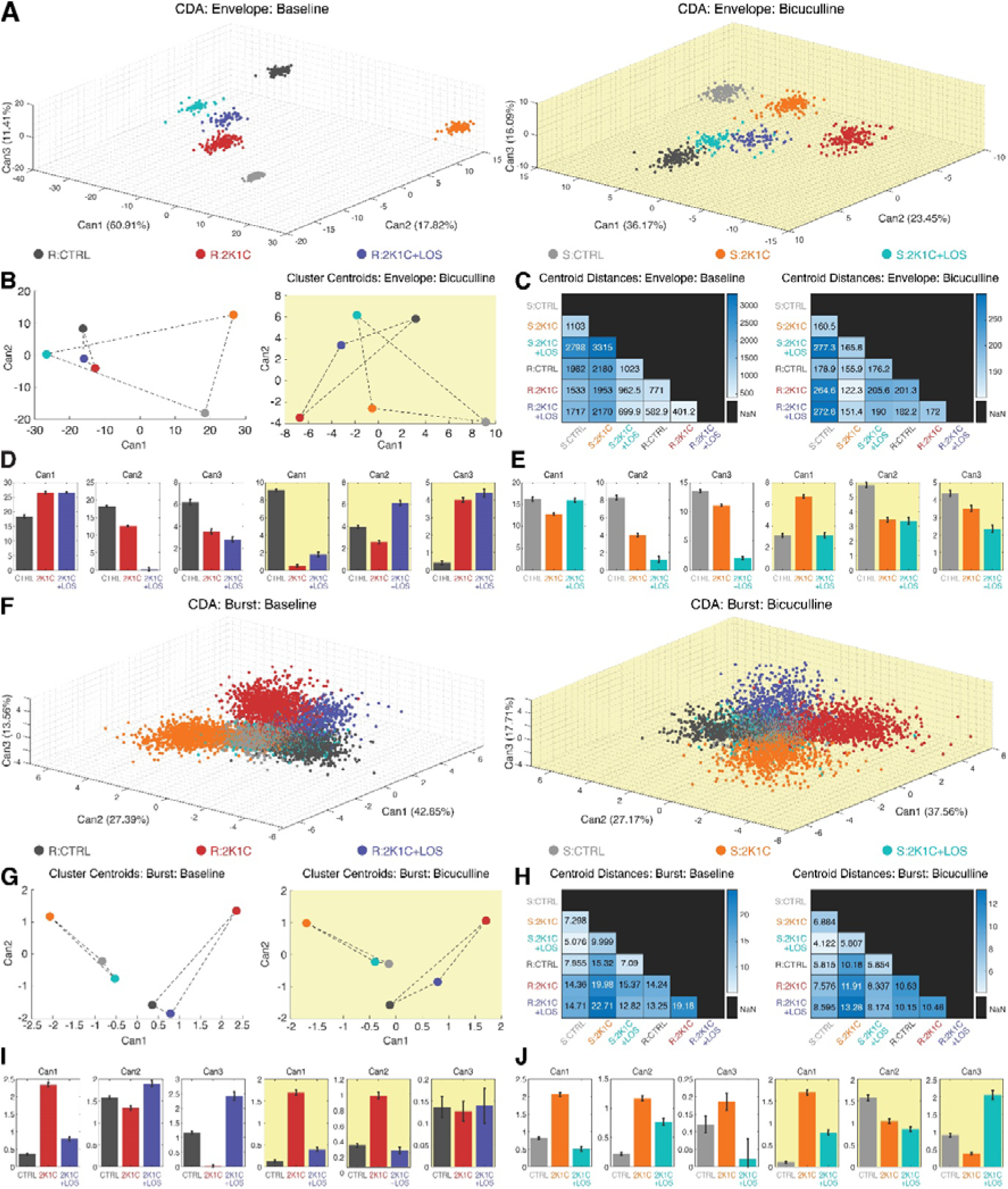
Canonical Discriminant Analysis (CDA) revealed specific frequency patterns for each nerve and each group in each condition, in both envelope and burst oscillations. **(A, F)** 3D scatter plot of the three first canonical variables from the clustering analysis using Canonical Discriminant Analysis (CDA) of both rSNA and sSNA envelope oscillations (A) and high frequency burst oscillations (F) at baseline (left, white background) and after bicuculline injection (right, yellow background). It’s possible to notice the clusters separation at visual inspection. **(B, G)** Clusters’ centroid plotted using the first and second canonical variables (Can1 and Can2), that discriminated between each of the groups. **(C, H)** Matrix of the centroid distances for each pair of groups’ clusters from baseline (left) and after bicuculline injection (right). **(D, E, I, J)** Confidence intervals of the first three canonical variables for the rSNA (D, I) and sSNA (E, J) at the baseline (left graphs, white background) and after bicuculline injection (right graphs, yellow background). (D, E, I, J) Values plotted as mean ± SEM. *p<0.05 (One-way ANOVA followed by Bonferroni’s post-hoc test). **(A-J)** Dark gray: Renal nerve, CTRL group; Dark blue: Renal nerve, 2K1C group; Red: Renal nerve, 2K1C+LOS group; Light gray: Splanchnic nerve, CTRL group; Orange: Splanchnic nerve, 2K1C group; Turquoise: Splanchnic nerve, 2K1C+LOS group.

Regarding the envelope oscillations analysis, all clusters are clearly separated in the three-dimensional space and statistically different at the baseline activity (Fig. 4A, left panel). This statistical trend can also be seen after the administration of bicuculline (Fig. 4A, right panel), although the clusters are spatially closer at visual inspection compared to the baseline condition, suggesting that the increased excitation in the PVN tends to reduce the differences of oscillatory patterns among groups and conditions.

This observation is supported by the analysis of clusters’ centroids distances. The centroid position is calculated by the average coordinates of all points that form a cluster. Since each cluster represents the spectral activity distribution for a specific group and nerve, the distance between two centroids reflects the degree of similarity between their rhythmic patterns. In other words, changes in the neural activity pattern of a given group/nerve can be detected by analyzing the distances between clusters in the frequency space. After bicuculline injection in the PVN, the centroids’ relative distance values dropped down up to 20 times (Fig. 4C). Interestingly, the envelope oscillating patterns of the renal nerve are more similar among the three conditions than the splanchnic nerve at the baseline (Fig. 4B). It’s noteworthy that the 2K1C+LOS group (blue dot) is almost equidistant from the CTRL group and the 2K1C group in both nerves and conditions, except for the renal nerve at baseline, suggesting that it induces an oscillatory effect totally different from the other two groups. Additionally, figures 4D (renal) and 4E (splanchnic) show the confidence interval comparisons among the canonical variables averages of each nerve for the three groups with and without bicuculline.

The same set of analysis was carried out considering the whole frequency range to include the high frequency oscillations of the bursting activity. The cluster analysis with the first three canonical variables from the CDA included all five animals from each group and both nerves. At visual inspection, it can be seen that at baseline the parameters are still well clustered (Fig. 4F, left panel), although not as separated as for the envelope oscillations (Fig. 4A, left panel). Bicuculline injection at the PVN also seems to have decreased the distance among clusters (Fig. 4F, right panel), although apparently to a less extent than for the envelope oscillations (Fig. 4A, right panel).

Analysis of the clusters’ centroid distances, however, revealed a linear separation among groups in both nerves and both conditions (with and without bicuculline), which can be noticed by the absence of intersection between the triangles formed by the lines joining the groups of each nerve (Fig. 4G). Furthermore, it’s noteworthy that the centroids of CTRL and 2K1C+LOS groups are remarkably closer in the high frequency analysis than in the envelope oscillations in both conditions, which is true even if we consider both nerves together (Fig. 4G). In other words, the four groups (R:CTRL, R:2K1C+LOS, S:CTRL and S:2K1C+LOS) share relatively similar oscillation patterns that are, in turn, rather distinct from the hypertensive groups, which is also supported by the statistical effects among groups in the confidence interval comparisons (Fig. 4I and Fig. 4J). Thus, the blockade of AT1 receptors by losartan seems to act differently over the high frequency bursts and the envelope oscillations, with a more prominent effect in recovering a normotensive pattern only in the high frequency bursts.

Additionally, the bicuculline injection in the PVN has a much milder effect over the high frequency bursts in decreasing the distances among clusters’ centroids when compared to the envelope oscillations (Fig. 4H). Hence, although the bicuculline administration in the PVN seems to have a general effect of increased oscillatory synchronization, this effect is more prominent over the envelope oscillations. The hyperexcitability at the PVN driven by bicuculline injection seems to entrain the structural properties of the circuits of each pathway with some kind of rhythmic marker, which could explain the rhythmic signature of each one.

## 4. DISCUSSION

In the present study we investigated the role of the PVN on the vasomotor sympathetic activity regulation by disturbing its inhibitory tonus with the injection of the GABAergic antagonist bicuculline, recording the renal and splanchnic nerves activity and comparing baseline and bicuculline conditions. Subjects were control and Goldblatt hypertensive rats, with one group of the latter being systemically treated with losartan, an antagonist of AT1 receptors with known antihypertensive effects. To the best of our knowledge, this is the first categorical demonstration that spectral parameters of the sympathetic vasomotor activity carry specific information to different territories that can be distinguished in both healthy and pathological conditions.

Our findings support the notion that the mean global power is a rather limited parameter to distinguish among the sympathetic nerve activity of control, hypertensive and hypertensive treated animals. Mean global power was only effective to distinguish between activity above and below 60 Hz (Fig. 3J and 3L, left panels) and between conditions (baseline and bicuculline) in the range below 4 Hz (Fig. 3C and 3E, left panels). In other words, mean global power is unable to yield refined information that distinguishes between groups. These results also support the idea that frequency and amplitude are independently modulated by the PVN in the sympathetic vasomotor activity.

The postganglionic sympathetic nerve activity is organized in bursts of high frequency action potentials between 100 Hz and 150 Hz (Fig. 2G and 2H, upper panels) that repeat in a periodic manner generating a rhythmic pattern of oscillatory steady state activity (Barman, 2020). The periodic pattern itself generates a virtual envelope frequency between 7 Hz and 10 Hz (Fig. 2A and 2B, upper panels). We therefore adopted an analysis approach dividing the recorded signal in two encoding mechanisms: High frequency bursts and low frequency envelope oscillations (Fig. 1C).

Using the shapes of the PSDs of the recorded signals (and therefore their spectral distribution) as the main feature for the CDA analysis, we found a highly clustered pattern of spectral organization among groups and nerves for the envelope oscillations at baseline (Fig. 4A, left panel), suggesting that each nerve has a highly specific spectral composition either under normal, hypertensive or treated conditions.

Since each cluster represents the distribution of spectral activity across different frequencies for a specific physiological condition and nerve, changes in the distances between these clusters indicate direct changes in their rhythmic patterns. In other words, alterations in the patterns of neural activity can be observed by analyzing the distances between clusters in the frequency space.

Bicuculline injection into the PVN dramatically decreased the distance among clusters’ centroids for the envelope oscillations, but not enough to bring them together (Fig. 4C). Since the clusters represent the nerve activity’s spectral features, this result means that the bicuculline injection increased the similarity of the rhythmic patterns among groups and nerves. It also changed the power distribution over the frequency range, flattening the activity above 2 Hz and concentrating the power spectrum in a typical oscillation pattern around 1.5 Hz (Fig. 4A and 4B).

Although not as separated as in the envelope oscillations, the clusters generated for the high frequencies by the same analysis also revealed distinct patterns of spectral organization, especially for frequencies above 60 Hz. The bicuculline microinjection into the PVN had a much softer effect in decreasing the clusters’ centroid distances than it had in the envelope oscillations.

Hence, these results suggest that the GABAergic neurons of the PVN play an important role in the temporal and spectral organization of the postganglionic sympathetic nerves bursting activity and in the differentiation of the spectral features coded in the reciprocal communication between them and the CNS. The absence of inhibitory modulation from the PVN has a milder effect over the higher frequencies but tends to converge the slow envelope oscillations of all nerves and groups into more similar spectral patterns. The high frequency activity is therefore probably generated at the structures other than the PVN, most likely local circuits. Nevertheless, they seem to be temporally organized by the slower envelope oscillations. However, once the patterns are still highly distinct after bicuculline injection, it’s highly unlikely that the PVN is the only structure generating the envelope frequency to distinct target territories. On the contrary, the spectral and temporal pattern of the slow envelope oscillations might be the result of the integrated activity of the PVN and other nuclei through some cross-frequency mechanism. One of the main types of these mechanisms is the phase-amplitude coupling, in which the amplitude of a high frequency oscillation is coupled to the phase of a low frequency oscillation, a wave morphology that resembles the bursting pattern of the renal and splanchnic nerves. phase-amplitude coupling has been extensively described in the CNS as the amplitude of a high frequency oscillation, such as a gamma (30-100 Hz), being coupled to the phase of a low frequency oscillation, such as delta (1-4 Hz) or theta (4-8 Hz) (Lisman and Jensen, 2013). Theta-Gamma and Delta-Gamma coupling have been functionally associated with the long range communication between hippocampus and prefrontal cortex respectively in declarative memory tasks (Colgin, 2016; Fell and Axmacher, 2011; Lisman and Jensen, 2013; Tort et al., 2009) and working memory tasks (Gągol et al., 2018; Lara and Wallis, 2015), suggesting that this mechanism is involved in the formation and retrieval of memory traces.

One possible explanation for this mechanism is that the slow oscillations provide traveling waves of excitability at whose peaks action potentials are more likely to happen. Therefore, by creating periodic windows of firing opportunity, slow oscillations would work as a probabilistic mechanism of temporal organization of fast spiking local circuits across long range networks (Buzsaki, 2006; Fries, 2005; Tal et al., 2020). Phase coherence between the activity of two given circuits and slow wave oscillations would be the key feature for integration. Theta oscillations underlie the integration of a network composed by the lateral amygdala, the infralimbic area of the medial prefrontal cortex and the CA1 area of the hippocampus during retrieval of conditioned fear in a contextual fear conditioning paradigm (Lesting et al., 2011). During extinction, coupling between LA and CA1 decreases, but this process can be delayed if both sites are electrically stimulated with synched in phase theta bursts, but not with a phase shift of 180º between sites (Lesting et al., 2011). This result shows how crucial the temporal organization is for the functional long distance coupling of different areas during a specific task. We hypothesize that a similar mechanism underlies the functional integration of brainstem, basal forebrain and spinal cord nuclei (such as the PVN, the rostral ventrolateral medulla and the intermediate lateral horn) with feedback information coming from sensors as baroreceptors. This network would control the differential activation of sympathetic nerves to specific target territories through dynamic organization of bits of information in phase with theta and/or delta bursts.

The blockade of the AT1 receptors via systemic treatment with losartan did not fully restore the baseline levels of autonomic symptoms and vasomotor activity in the Goldblatt hypertensive rats (Fig. 2). Notwithstanding, AT1 receptors somehow modulate the power spectral distribution in both nerves, once there were significant differences between the spectral parameters of the group 2K1C+LOS in relation to the others in almost all comparisons in the CDA analysis (Fig 4). The fact that systemic treatment with losartan did not restore the MAP levels suggests that the changes in the sympathetic vasomotor activity triggered by the PVN’s hyperexcitability are not dependent on blood pressure, although we don’t know how long such alterations persist after the normalization of blood pressure levels by losartan treatment.

The losartan effect on the envelope oscillations, however, followed quite different patterns between renal and splanchnic nerves, especially for the envelope oscillations (Fig. 4A to 4E). This result can be explained by the fact that the sympathetic vasomotor activity is topographically controlled through differential coding by the CNS, i.e. the activity pattern to the kidneys can have distinct intrinsic parameters than the splanchnic territory and vice-versa. In addition, with the fact that frequency and amplitude of vasomotor activity can be also independently controlled by the brain, it’s not surprising that the vasomotor responses induced by the losartan treatment were different for the renal and splanchnic territories mainly regarding the envelope oscillations, but not much to the high frequency activity (McAllen, R. M., and Malpas, 1997). Nevertheless, it should be noted that the losartan treatment in the present study took place for only one week and it would be important to investigate whether the mechanisms generating the sympathetic vasomotor activity would be modified by a longer treatment.

In summary, our results strongly suggest that frequency-encoded patterns are part of the mechanisms underlying vasomotor sympathetic activation in both normal and hypertensive conditions and that these patterns are modulated by the inhibitory tonus of the PVN. Although it was not possible to dissociate the intrinsic effects of the virtual envelope oscillations over the high frequency bursts, the isolated analysis of the envelope oscillations suggested a possible amplitude-frequency coupling effect. To better understand its dynamics and role in the encoding and transmission of information in each nerve and condition, further analyses are still needed.

Our results will hopefully help to improve our understanding of the intricate mechanisms that control the sympathetic vasomotor activity and the genesis and maintenance of hypertension and renal failure. In addition, we hope that the signal analysis approach used in this study might contribute to the progress of the field with a new set of powerful tools and techniques.

## Data availability statement

The original contributions presented in this study are included in the article material, further inquiries can be directed to the corresponding authors.

## Authors contribution

Conceptualization: R.R.C., M.I.O.M., C.S.S., and J.F. Project coordination and funding: R.R.C. Project supervision: J.F. and R.R.C. Data acquisition: M.I.O.M. Figure preparation: C.S.S. Data analysis: J.F. Writing: C.S.S., J.F., R.R.C. and M.I.O.M. All authors read and reviewed the manuscript.

